# AP2XII-2 coordinates transcriptional repression for *Toxoplasma gondii* sexual commitment

**DOI:** 10.1101/2022.11.29.518462

**Authors:** Sandeep Srivastava, Michael J. Holmes, Michael W. White, William J. Sullivan

## Abstract

*Toxoplasma gondii* is a widespread protozoan parasite that has significant impact on human and veterinary health. The parasite undergoes a complex life cycle involving multiple hosts and developmental stages. How *Toxoplasma* transitions between life cycle stages is poorly understood, yet central to controlling transmission. Of particular neglect are the factors that contribute to its sexual development, which takes place exclusively in feline intestines. While epigenetic repressors have been shown to play an important role in silencing spurious gene expression of sexually committed parasites, the specific factors that recruit this generalized machinery to the appropriate genes remains largely unexplored. Here, we establish that a member of the AP2 transcription factor family, AP2XII-2, is targeted to genomic loci associated with sexually committed parasites along with the epigenetic regulators of transcriptional silencing, HDAC3 and MORC. Despite widespread association with gene promoters, AP2XII-2 is required for silencing of relatively few genes. Using CUT&Tag methodology, we identify two major genes associated with sexual development downstream of AP2XII-2 control, AP2X-10 and the amino acid hydroxylase AAH1. Our findings show that AP2XII-2 is a key contributor to the gene regulatory pathways modulating *Toxoplasma* sexual development.

**IMPORTANCE:** *Toxoplasma gondii* is a parasite that undergoes its sexual stage exclusively in feline intestines, making cats a major source of transmission. A better understanding of the proteins controlling the parasite’s life cycle stage transitions is needed for the development of new therapies aimed to treat toxoplasmosis and transmission of the infection. Genes that regulate the sexual stages need to be turned on and off at the appropriate times, activities that are mediated by specific transcription factors that recruit general machinery to silence or activate gene expression. In this study, we identify a transcription factor called AP2XII-2 as being important for repression of a subset of sexual stage genes, including a sexual stage-specific AP2 factor (AP2X-10) and a protein (AAH1) required to construct the infectious oocysts expelled by infected cats.

## INTRODUCTION

The ubiquitous parasite *Toxoplasma gondii* (phylum Apicomplexa) infects warm-blooded animals the world over, including approximately one in three people (1). After an initial acute phase of infection by tachyzoites, the parasites redistribute within the solid organs of the infected host and reside as bradyzoites in latent tissue cysts that appear impervious to the immune response and currently available treatments (1). Persistence within the host makes *Toxoplasma* an opportunistic infection in those with compromised immune systems (1). Upon immune suppression, latent bradyzoite stage parasites reconvert into rapidly growing tachyzoites, causing severe tissue destruction that can threaten the life of the infected individual (1).

Given the importance of interconversion between tachyzoites and bradyzoites to disease progression, the molecular mechanisms governing this stage transition have received a great deal of attention (2). The conversion of tachyzoites to bradyzoites requires extensive reprogramming of gene expression at multiple levels, directed in part by chromatin remodeling histone acetyltransferases and deacetylases that are recruited to specific gene loci through their interactions with sequence-specific transcription factors (3, 4). In support of this, members of the AP2 transcription factor family have been found to associate with a cadre of histone modifying complexes including GCN5b/ADA2a and HDAC3/MORC (5, 6). AP2 transcription factors harbor a plant-like DNA-binding domain and have been implicated in contributing to the regulation of gene expression in apicomplexan parasites (7).

In contrast, our understanding of the molecular mechanisms governing the rewiring of gene expression during *Toxoplasma* sexual development is far less developed, largely due to a lack of model systems. This is despite evidence showing that ingestion of the ultimate product of *Toxoplasma* sexual reproduction, sporulated oocysts, accounts for a substantial portion of worldwide human infections and mass outbreaks of acute toxoplasmosis (8, 9).

The early stages of *Toxoplasma* sexual development occur in the intestinal lining of cats. Upon ingestion by the feline, the parasites invade the intestinal enteroepithelial cells and undergo several rounds of schizogonic replication to produce merozoites (10). Successive rounds of schizogony have been subcategorized into types A-E of the enteroepithelial stage. These pre-sexual stages then undergo gametocytogenesis, ultimately producing motile male microgametes and intracellular female macrogametes. After fertilization, up to a billion immature oocysts are expelled with the cat’s feces (11). Upon exposure to the air, the oocysts sporulate and become capable of transmitting infection (10).

It has recently been shown that transcriptional repression mediated by the HDAC3/MORC complex plays an important role in preventing the aberrant expression of genes normally restricted to the sexual stages in felids (6). The HDAC3/MORC complex associates with at least 11 AP2 family transcriptional regulators (6, 12), suggesting that these factors mediate targeted silencing of felid-specific developmental gene expression, as proposed for bradyzoite formation (4).

We initially found AP2XII-2 to be in association with AP2IX-4, a transcriptional repressor that helps coordinate bradyzoite formation (12, 13). We then reported that AP2XII-2 is required for proper progression of parasites through the S-phase of the tachyzoite cell cycle and that knockdown of AP2XII-2 increased frequency of bradyzoite formation *in vitro* (12). We and others have also reported the association between the HDAC3/MORC complex and AP2XII-2 (6, 12), suggesting that this AP2 family member controls the transcriptional repression of certain genes. However, the specific genes regulated by AP2XII-2 remained unexplored.

To better understand the role of AP2XII-2 in *Toxoplasma* biology, we profiled the genome-wide chromatin-binding sites of AP2XII-2. By adopting Cleavage Under Targets and Tagmentation (CUT&Tag) for use in *Toxoplasma,* we found find a high degree of overlap between AP2XII-2 and HDAC3/MORC target genes. Using a transcriptomics approach, we also found that AP2XII-2 controls the expression of a subset of HDAC3/MORC-target genes, including some that are restricted to the sexually-committed developmental stages such as AP2X-10 and the aromatic amino acid hydroxylase, AAH1. Our results indicate that AP2XII-2 helps coordinate recruitment of the HDAC3/MORC complex to specific gene loci to repress developmentally-controlled genes.

## RESULTS AND DISCUSSION

### AP2XII-2 shows widespread occupancy at gene promoters and high degree of overlap with the HDAC3/MORC complex

To identify genes that are direct targets of AP2XII-2 in tachyzoites, we applied the CUT&Tag method (14) to profile the genomic loci bound by the factor. To pulldown AP2XII-2 for CUT&Tag, we took advantage of a parasite clone we made previously in ME49 strain that expresses an HA-tagged version of the endogenous protein (12). Results showed that AP2XII-2-associated regions are widespread across the parasite genome, displaying a total of 5,527 peaks associated with 3,939 genes (Table S1; Fig. 1A). Consistent with its putative role as a transcriptional regulator, AP2XII-2 is located near the transcriptional start site of protein coding genes (Fig. 1B).

**Figure 1.**
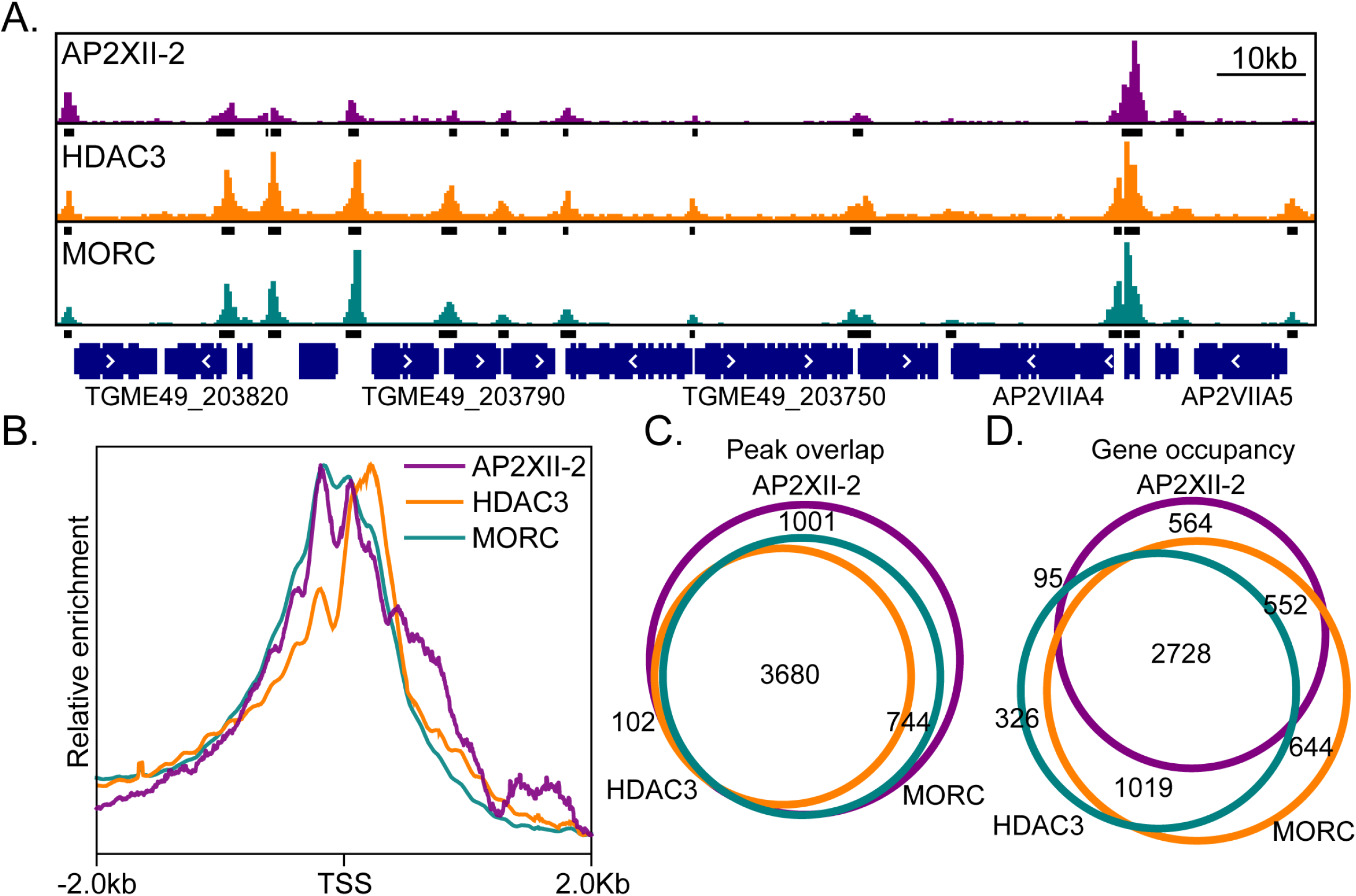
AP2XII-2 genome occupancy significantly overlaps with HDAC3 and MORC. A) AP2XII-2 genome occupancy along a segment of chromosome VIIA reveals its association at the promoters of several genes. Statistically enriched segments are shown as black bars directly under AP2XII-2 enriched peaks (purple). A similar distribution and enrichment of HDAC3 and MORC is seen in the orange and teal tracks, respectively. B) Genome-wide analysis reveals each member of the transcriptional repression complex associates with the transcriptional start sites (TSS) of genes. C) Venn diagram indicating that the majority of AP2XII-2 enriched peaks fall within those identified for HDAC3 and MORC. D) Venn diagram indicating that the majority genes occupied by AP2XII-2 are also occupied by HDAC3 and MORC.

Previous reports have demonstrated that AP2XII-2 is a member of the multi-component HDAC3/MORC complex, which silences genes by inducing heterochromatin formation (6, 12). To corroborate this association, we re-analyzed the existing HDAC3 and MORC ChIPseq data (6) to determine the overlap between HDAC3, MORC, and AP2XII-2 localization genome-wide (Table S1). Like AP2XII-2, both HDAC3 and MORC are enriched at the transcriptional start site of protein coding genes (Fig. 1B). Just over 80% of AP2XII-2 enriched regions overlapped with HDAC3 and/or MORC enriched peaks (Fig. 1C). Additionally, over 85% of genes associated with AP2XII-2 were also bound by HDAC3 and/or MORC with a clear plurality of all identified genes bound by the three proteins (Fig. 1D). Given that AP2 domains bind distinct DNA motifs and that HDAC3 and MORC lack DNA binding domains (15, 16), combined with the previous observation that they operate within a shared complex (6, 12), the high degree of coordination between AP2XII-2, HDAC3, and MORC binding sites strongly suggests that AP2XII-2 is responsible for recruiting the HDAC3/MORC complex to repress specific genes.

### Loss of AP2XII-2 disrupts a small subset of protein coding transcripts in tachyzoites

Given that AP2XII-2 associates with a large number of gene promoters and likely directs transcriptional silencing through the HDAC3/MORC complex, we wanted to assess the consequences of AP2XII-2 depletion on transcriptional regulation. To maximize the robustness of this experiment, we compared the response to AP2XII-2 depletion in both type I (RH strain) and type II (ME49 strain) backgrounds using parasites we made in a previous study (12). These parasites express endogenous AP2XII-2 tagged at its C-terminus with a dual tag comprised of an HA epitope and an auxin-inducible degron (AID). TheAP2XII-2^AID-HA^ protein in both strains was degraded to undetectable levels within 24 hours of adding 500 μM indole-3-acetic acid (IAA) to the culture media (Fig. 2A).

**Figure 2.**
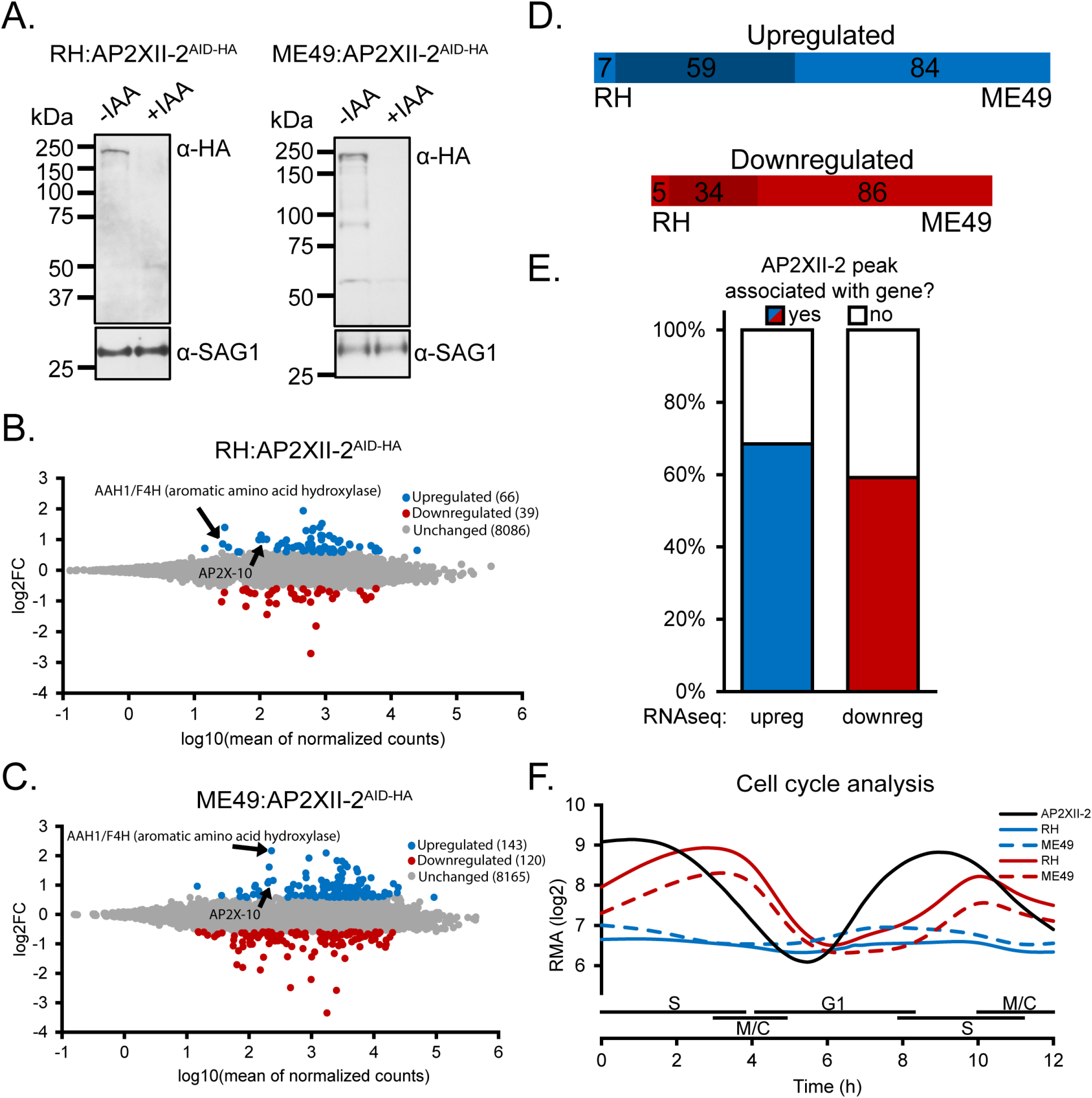
AP2XII-2 depletion leads to relatively little gene dysregulation. A) Western blots confirming IAA-dependent downregulation of AP2XII-2^AID-HA^ protein in RH^TIR1^ or ME49^TIR1^ parasites after 24 hours incubation with 500 μM IAA in culture media. Blots were probed with anti-SAG1 as a loading control. B-C) MA plot of differential gene expression analysis of AP2XII-2^AID-HA^-depleted in RH^TIR1^ (B) or ME49^TIR1^ (C) parasites 24 hours after IAA addition. Significantly upregulated genes are colored in blue, downregulated genes in red. Genes of interest for further analysis are noted. D) Diagrams showing overlap of significantly upregulated and downregulated genes in ME49 and RH genetic backgrounds. E) Analysis of AP2XII-2 gene occupancy and directionality of dysregulation that occurs upon AP2XII-2 depletion. F) Meta cell cycle expression analysis of significantly upregulated (blue) and downregulated (red) gene sets that respond to AP2XII-2 depletion. The cell cycle profile of AP2XII-2 is shown in black as a reference.

Using these conditions, comparative RNA sequencing (RNAseq) analyses revealed that the loss of AP2XII-2 in tachyzoites led to dysregulation of mRNA levels for a modest number of transcripts in both the RH and ME49 genetic backgrounds (Table S2). In RH parasites lacking AP2XII-2, 66 genes were significantly upregulated and 39 genes were downregulated at least 1.5 fold (Fig. 2B). Similarly, in ME49 parasites, 143 genes were significantly upregulated and 120 genes were downregulated 1.5 fold following depletion of AP2XII-2 (Fig. 2C). The majority of the dysregulated genes in RH strain were also dysregulated in ME49 strain, indicating a high degree of concurrence between strains (Fig. 2D). Surprisingly, there was no correlation between the direction of transcriptional dysregulation and AP2XII-2 gene occupancy as determined by CUT&Tag (Fig. 2E). This observation likely speaks to the complexity of gene regulation in *Toxoplasma*, which can involve multiple AP2 factors and chromatin remodeling machinery. Depletion of a single factor like AP2XII-2 may not be sufficient to significantly disrupt expression of most genes in the network. Since AP2XII-2 was also found to interact with AP2IX-4 (12), our observations here may point to a model where AP2 family members cooperatively bind for efficient HDAC3/MORC recruitment to specific gene loci.

Since AP2XII-2 is a cell cycle regulated factor with expression peaking during the S/M phases (12, 17), we assessed whether the dysregulated genes were also subject to cyclic control. Using the cell cycle microarray data available on ToxoDB.org (17, 18), we determined the average cell cycle profiles for the genes that were up- or downregulated in response to AP2XII-2 depletion and compared them to the cell cycle pattern exhibited by AP2XII-2 (Fig. 2F). Downregulated genes tended to be cell cycle regulated with a valley of expression that spans mid-G1 phase, in line with our previous observation that AP2XII-2 depletion delays the cell cycle after centrosome duplication (12). The expanded G1 phase occupancy induced by AP2XII-2 depletion may point to a linkage between transcriptional repression of AP2XII-2 target genes and the cell cycle. Despite not displaying cell cycle regulation, the upregulated genes were expressed at lower robust multichip average (RMA) expression values. The relatively low abundance of these genes in tachyzoites could indicate that they are normally subjected to transcriptional repression. Given the association of AP2XII-2 with the HDAC3/MORC transcriptional repression complex (6, 12), we suspected that these upregulated genes could be subject to regulation by AP2XII-2 via recruitment of HDAC3/MORC.

### Loss of AP2XII-2 increases expression of genes normally restricted to latent and sexually committed parasites

To identify genes that are most likely to be direct targets of the AP2XII-2-directed HDAC3/MORC complex, we looked for genes that were upregulated in 1) either ME49 or RH strain parasites upon AP2XII-2 depletion, 2) after 24 hours of MORC depletion (Fig. S1A), and 3) after 18 hours incubation with the HDAC3 inhibitor FR235222 (Fig. S1B). These latter two criteria were assessed using publicly available data and were also generated from a type II parasite strain (Table S3) (6). In addition, we verified that each gene’s promoter was occupied by AP2XII-2, HDAC3, and MORC. Five gene accession entries met this stringent filtering criteria (Table 1).

**Table 1.** Genes occupied by AP2XII-2, HDAC3, and MORC that are consistently upregulated upon their depletion or inhibition.

| Gene ID | Product Description | AP2XII-2 depletion |  | MORC<sup>a</sup> | HDAC3<sup>a</sup> |
| --- | --- | --- | --- | --- | --- |
|  |  | ME49<sup>a</sup> | RH<sup>a</sup> |  |  |
| TGME49_212710 | phenylalanine-4-hydroxylase (F4H) | 2.18 | 0.86 | 4.29 | 3.74 |
| TGME49_287510 | aromatic amino acid hydrolase (AAH1) | 0.58 | 1.40 | 4.91 | 3.22 |
| TGME49_318750 | deoxyribose-phosphate aldolase (TgDPA) | 1.82 | 1.30 | 1.39 | 1.40 |
| TGME49_215340 | AP2X-10 (fragment) | 1.16 | 1.01 | 5.19 | 3.57 |
| TGME49_215343 | hypothetical protein (AP2X-10 fragment) | 1.13 | 1.00 | 5.27 | 3.45 |

<sup>a</sup> log<sub>2</sub>FC values

The auxin-induced depletion of MORC as well as inhibition of HDAC3 in tachyzoites induced upregulation of bradyzoite-specific genes as well as genes associated with sexual stages (6, 19). Notably, when assessing the degree of dysregulation for the genes outlined in Table 1, the effect was more pronounced for MORC and HDAC3 than for AP2XII-2 depletion. This observation is consistent with a model whereby multiple AP2 family members cooperatively recruit the HDAC3/MORC complex to genomic loci.

It has been proposed that parasites must pass through the bradyzoite stage before initiating sexual commitment (6, 20). Our analysis (Table I) identified a single bradyzoite-specific gene, TgDPA (21), to be reliably upregulated under all three treatments that were assessed, strongly suggesting that its expression is controlled by an AP2XII-2-directed HDAC3/MORC complex. The role that TgDPA plays in bradyzoite biology and their potential commitment to sexual development remain to be elucidated.

Interestingly, the AP2 family member AP2X-10, whose current fragmented gene annotation incorrectly assigns the gene to both TGME49_215340 and TGME49_215343, is upregulated when either AP2XII-2, MORC, or HDAC3 is impaired independent of parasite strain (Table 1). AP2X-10 expression has been noted to be restricted to the enteroepithelial and oocyst stages; it is the most abundant of all AP2 factors in oocysts, being expressed at the 96^th^ percentile (22–24). Seventeen AP2 family members have been proposed to act downstream of the HDAC3/MORC complex to regulate various developmental trajectories, with AP2X-10 predicted to guide micro- and/or macrogamete development (6). Our present study is consistent with AP2XII-2 coordinating the repression of AP2X-10, placing the former above AP2X-10 in a transcriptional regulatory cascade that contributes to the coordination of *Toxoplasma* sexual development. Future studies to determine the genes directly controlled by AP2X-10 and clarify its role in sexual development are warranted.

### AAH family gene organization

Two aromatic amino acid hydroxylase enzymes, AAH1 and F4H, are not only counted amongst the highest upregulated genes upon AP2XII-2, MORC, and HDAC3 inhibition (Table 1), but have also been reported to be important for oocyst maturation (25). Interestingly, we found that F4H was upregulated in ME49 but not RH strain, AAH1 showed the opposite trend (Table 1). These observations prompted us to further examine the entire aromatic amino acid hydroxylase family in *Toxoplasma*, which consists of three annotated members, TGME49_287510 (AAH1), TGME49_212740 (AAH2), and TGME49_212710 (F4H).

Previous reports have indicated that the current ME49 genome assembly available on ToxoDB.org has incorrectly fused a tandem repeat of the AAH2 gene (25, 26). Before proceeding with our analysis, we had to ensure the fidelity of the AAH2 gene sequence. To that end, we compared the ME49 genome assembly with the RH88 assembly since the former was generated using short read sequencing technology, which can have difficulty with repetitive regions (Fig. 3A). The RH88 genome was constructed based on long read sequencing technology, which is better at determining overall genome architecture (27). This comparison revealed that indeed two members of the AAH family should appear on chr V of the ME49 genome, spaced apart by unplaced contigs KE139705 and KE139818 (Fig. 3A).

**Figure 3.**
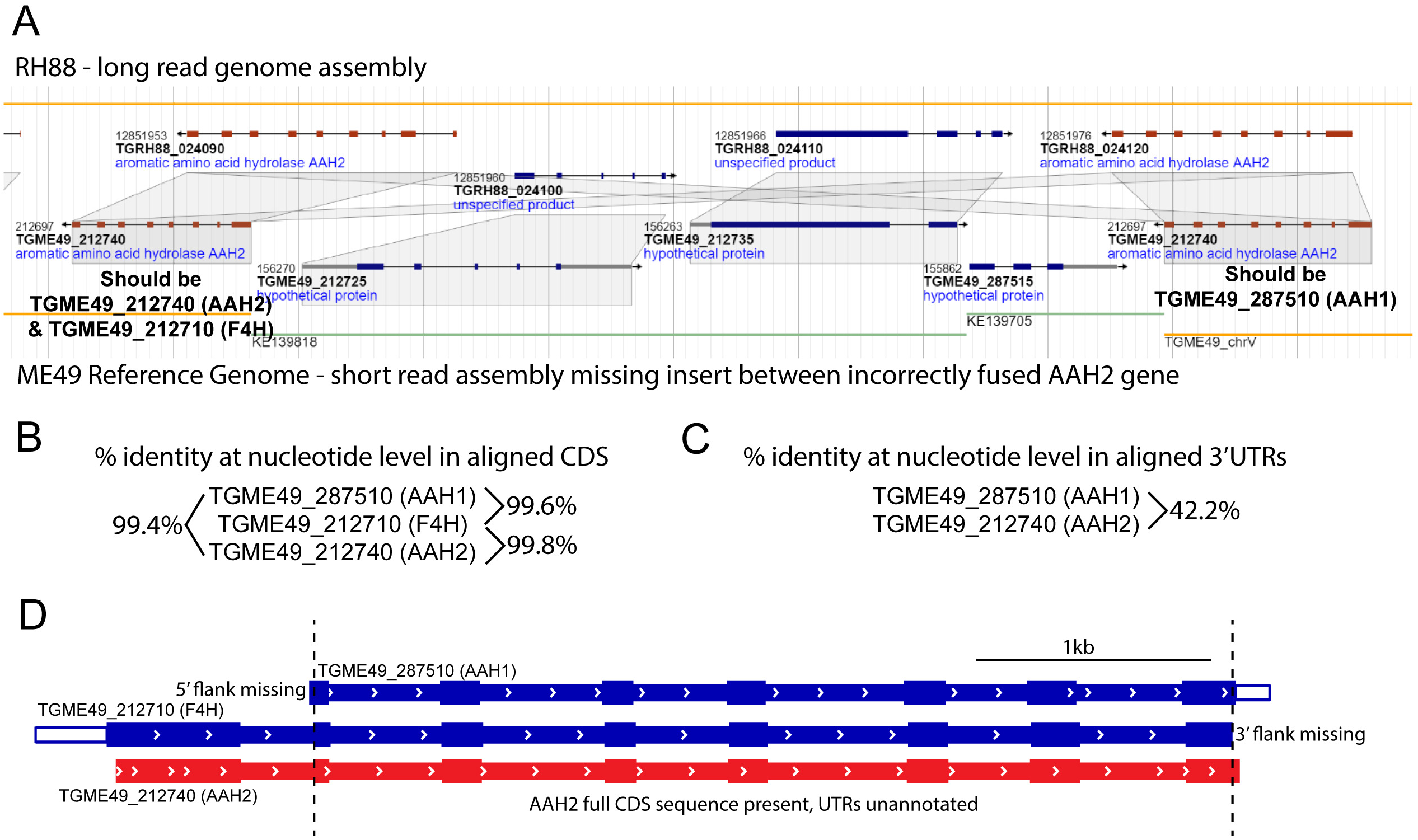
Disentanglement of the aromatic amino acid hydroxylase family of genes on chr V. A) Genomic alignment and syntenic analysis of RH88 and ME49 genomes available on ToxoDB.org reveals the correct positioning of unplaced contigs KE139705 and KE139818 into chr V that was missed due to incorrect fusion of AAH1/2 genes in the ME49 reference strain. Suggested changes to annotation of AAH1 and AAH2 are based on sequence alignments of 3’ flanking regions in both genes across both parasite strains. B) Multiple pairwise alignments of CDS regions that are present in each AAH gene fragment reveals a high degree of identity at the nucleotide level which obfuscates gene calling during differential gene expression analysis. C) Pairwise alignment of 3’ flanking regions reveals a clear way to discriminate both AAH1 and AAH2 genes for C-terminal tagging at endogenous loci. D) Outline of AAH-family gene fragments available in ME49 reference strain. Missing fragments on either 5’ or 3’ terminus are stated. Conserved CDS regions that were used for multiple sequence alignment are delimited by dotted line.

The three ME49 members of the AAH family share over 99.4% identity at the nucleotide level in their coding sequences (Fig. 3B), suggesting that the vast majority of RNAseq reads mapping to the CDS region would have been assigned to all AAH genes indiscriminately; this spurious read assignment may indicate that a putative strain-specific upregulation of AAH1 or F4H could occur by chance (Table 1). Given the high degree of sequence identity between each AAH family member, we examined their flanking sequences to differentiate them. AAH1 and AAH2 have unique 3’end flanking sequences (Fig. 3C), however the unplaced contig KE139818 abruptly ends, leaving the annotated F4H gene (TGME49_212710) without a 3’UTR sequence (Fig. 3D). Analysis of the unique regions flanking the 5’ and 3’ ends of the AAH family by sequence alignment comparison with the RH88 genome architecture allowed us to place AAH1 and AAH2 onto chr V and determine that F4H and AAH2 are likely the same gene (Fig. 3A). However, it should be noted that a previously conducted copy number variant analysis of the AAH family in *Toxoplasma* indicated that ME49 parasites harbor three copies of AAH genes whereas type I parasites encode only two (25). We were unable to map the location of the additional AAH family member with our strategy of comparing ME49 and RH genome architectures.

### Depletion of AP2XII-2 de-represses AAH1 but not AAH2

To distinguish which AAH family member(s) are upregulated upon AP2XII-2 depletion, we endogenously tagged AAH1 and AAH2 with MYC epitopes at their C-termini in ME49 AP2XII-2^AID-HA^ parasites. The tagging of these genes was confirmed by verifying successful recombination at each genetic locus by PCR analysis (Fig. S2). In untreated parasites, AAH1^MYC^ was not detectable by western blotting whereas AAH1^MYC^ appeared at the expected size of 62 kDa upon IAA-induced loss of AP2XII-2 (Fig. 4A). Expression of AAH2^MYC^ was not seen under either condition (Fig. 4B).

**Figure 4.**
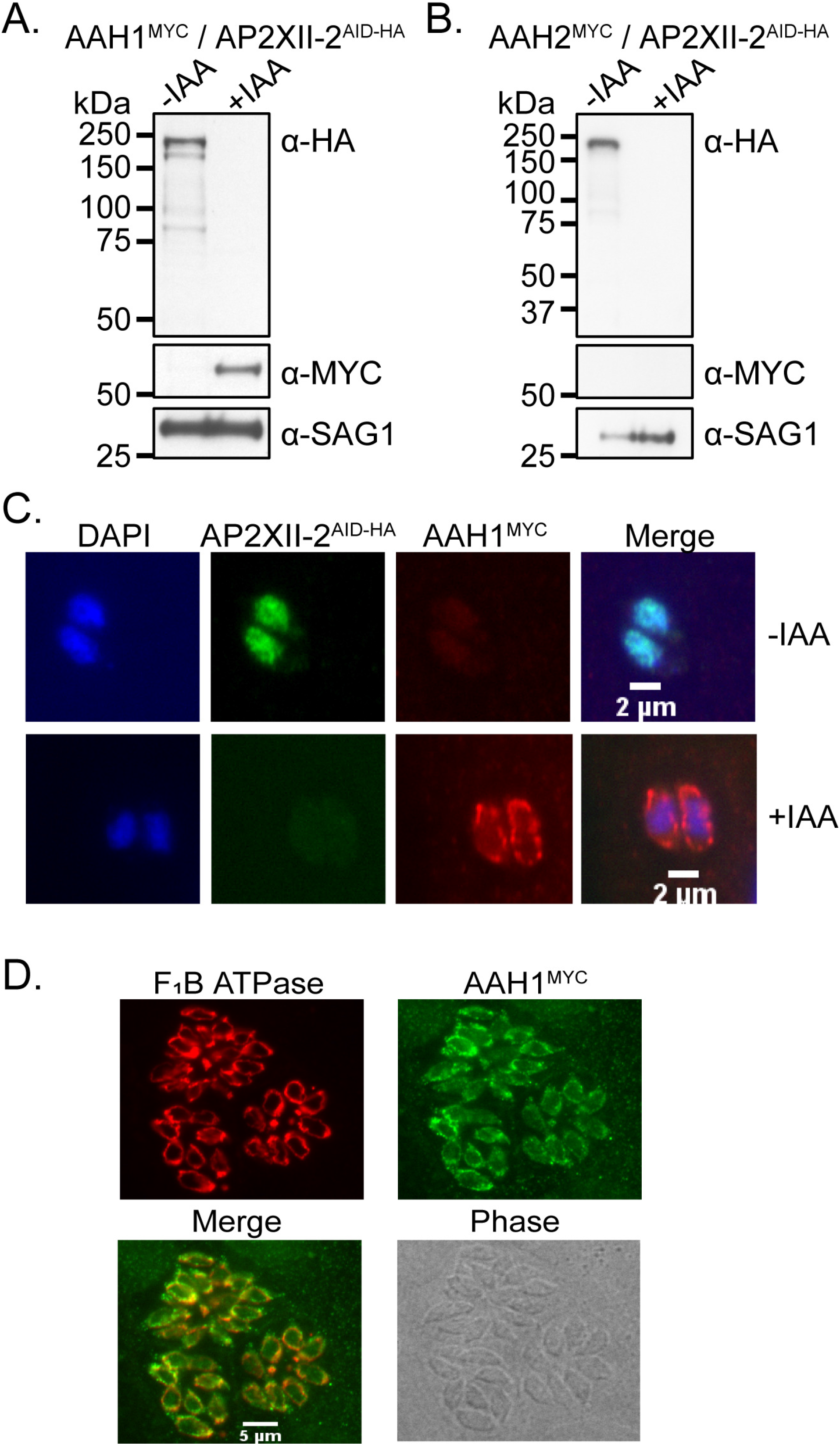
Depletion of AP2XII-2 causes de-repression of AAH1 gene expression. A-B) Western blots demonstrating that AAH1 (A) and not AAH2 (B) is expressed upon IAA-induced depletion of AP2XII-2. Both genes were endogenously tagged at their C-termini with MYC epitope tags. Blots were probed with anti-SAG1 as a loading control. C) AAH1 expression is observed in a concentric pattern around the nucleus upon AP2XII-2 depletion. D) AAH1 is expressed in the vicinity of the parasite mitochondrial marker F_1_BATPase.

Neither AAH family member is typically expressed in asexual parasites (26), suggesting that they are maintained in a transcriptionally repressed state in non-sexually committed parasites. Our results suggest that AAH genes may be regulated by different HDAC3/MORC-AP2 complexes, with AP2XII-2 directing the silencing of AAH1 but not AAH2. However, we cannot rule out the possibility that AAH1 and AAH2 have a parasite strain specific dependence on AP2XII-2. Since parasites lacking AAH1 have a severe defect in infection of the cat intestine and oocyst production (25), we propose that AP2XII-2 is a key factor required for completion of *Toxoplasma* sexual reproduction.

We next examined AAH1 expression by immunofluorescence assay (IFA) and found AAH1^MYC^ to be localized around the nuclei of tachyzoites only upon AP2XII-2 depletion (Fig. 4C). This staining pattern is reminiscent of the *Toxoplasma* mitochondrion. To further characterize AAH1^MYC^ localization, we co-stained AP2XII-2^AID-HA^ knockdown parasites with antibodies to the F_1_B ATPase mitochondrial marker (28) and the MYC epitope tag, which confirmed the co-localization of AAH1^MYC^ with the mitochondrion in tachyzoites (Fig. 4D).

The AAH family of enzymes is responsible for converting phenylalanine to tyrosine and tyrosine to 3,4-dihydroxy-L-phenylalanine (L-DOPA) (29). The later compound is a precursor for the formation of di-tyrosine protein crosslinks that are thought to play a critical structural role in *Toxoplasma* oocyst wall integrity and formation (25). Given the role that AAH family enzymes play in ensuring oocyst development and wall integrity, we were surprised to see AAH1 localized to the parasite mitochondrion. Interestingly, a previous report demonstrated that AAH2 overexpression in bradyzoites gave rise to a staining pattern consistent with mitochondrial localization (26), suggesting that the mitochondrion might act as a hub for AAH enzymatic activity. Future studies using bona fide sexually-committed parasites are needed to verify this possibility.

### Conclusion

We have determined that in addition to affecting tachyzoite cell cycle progression (12), AP2XII-2 also plays a role in coordinating *Toxoplasma* sexual development. This may indicate that AP2XII-2 plays different roles in different developmental stages or may suggest that the commitment to sexual development is a cell-cycle dependent phenomenon as has been demonstrated for the tachyzoite to bradyzoite transition (7, 30). Consistent with this model, either inhibition of HDAC3 or MORC depletion also stall cell cycle progression and lead to increased bradyzoite and sexual stage gene expression (6, 19), albeit to a greater extent than does AP2XII-2 depletion. Given the association of AP2XII-2 with AP2IX-4 (12), and the observation that AP2XII-2 depletion appears to cause a milder version of the HDAC3 and MORC phenotypes, we propose that AP2XII-2 is likely one of several AP2 family members that bind cooperatively to recruit the HDAC3/MORC to specific genomic loci.

On the other hand, the number of reports demonstrating that *Toxoplasma* studies investigating gene regulation using in vitro culture system lack the fidelity of in vivo parasite development (31–33). This is particularly relevant when examining the roles of AP2 family members in gene regulation (33). Notably, a previous report indicating that AP2IX-9 is a repressor of bradyzoite formation was later contradicted using a developmentally competent *Toxoplasma* models (15, 22, 33). Given that we have reported that AP2XII-2 impacts both cell cycle regulation (12) and the expression of sexually committed gene expression, we cannot rule out the possibility that mis-expression of AP2XII-2 may have resulted in ‘transcriptional confusion’ that impacted tachyzoite cell cycle dynamics. Examining the role that AP2XII-2 plays throughout the complete life cycle stages of *Toxoplasma* development is warranted.

The intrinsic factors responsible for regulating *Toxoplasma* sexual development have been understudied due to the lack of amenable systems. This is beginning to change with the pivotal discoveries of the upstream extrinsic signals required to initiate sexual development and the requirement of widespread transcriptional repression to coordinate the process (6, 34). Future work will undoubtedly focus on disentangling the gene regulatory networks responsible for initiating and managing *Toxoplasma* sexual development. Here we have begun unravelling one of the networks responsible for this process by placing the AP2XII-2-directed HDAC3/MORC complex above two genes that function in sexual stages, AP2X-10 and AAH1. In addition to utilizing a candidate-based approach as we have relied on for this study, the recent report highlighting inhibitors of AP2-family member activity may prove yet another tool for interrogating the role of AP2 factors in *Toxoplasma* developmental transitions (35).

## MATERIALS AND METHODS

### Parasite culture

Human fibroblast cells (HFFs, ATCC SCRC-104) were maintained in Dulbecco’s modified Eagle medium (DMEM; Corning) supplemented with 10% fetal bovine serum (FBS, R&D systems), 100 μg/ml streptomycin and 100 units/ml penicillin as described elsewhere (36). RH and ME49 parasites were cultured in DMEM supplemented with 1% and 5% heat inactivated FBS, respectively, 100 μg/ml streptomycin and 100 units/ml penicillin. Parasites were maintained in humidified incubator at 37°C and 5% CO_2_. Transfected and clonal strains were maintained in selection media supplemented with 1 μM pyrimethamine or a combination of 25 μg mycophenolic acid and 50 μg xanthine as appropriate. Control of AP2XII-2 expression was achieved by adding either 500 μM of indole-3-acetic acid or an equivalent amount of ethanol vehicle for 24 hours to the culture media before analysis.

### Generation endogenously tagged AAH1^MYC^ or AAH2^MYC^

The ME49-TIR1:AP2XII-2^AID-HA^ strain that we generated in a previous study (12) was used as the parental strain for generating dual-tagged AAH1^MYC^ and AAH2^MYC^ parasites. For AAH1, the guide RNA sequence from the pSAG1::CAS9-U6::sgUPRT plasmid (Addgene 54467) (37) was mutagenized to target the stop codon of AAH1. All primers and sequences are listed in supplement (Table S4). A donor repair template with regions of homology to the AAH1 CDS and 3’UTR was made to incorporate the MYC tag and HXGPRT cassette. Parasites were transfected with 25 μg of CAS9 plasmid and 25 μg of donor repair template, were cultured with 25 μg mycophenolic acid and 50 μg xanthine for 7-8 days and single clones were isolated by limiting dilution in 96 wells. Isolated single clones were confirmed for C-terminal tagging by PCR of their genomic DNA (Fig. S2)

To generate AAH2^Myc^ parasites, specific primers were used to clone ~2 kb of the AAH2 gene into the PacI site of the pLIC-3xMYC-HXGPRT plasmid. 50 μg of the final construct was linearized with NheI restriction endonuclease and transfected into the ME49-TIR1:AP2XII-2^AID-HA^ strain. Transgenic parasites were selected for 7-8 days under drug pressure and subjected to limiting dilution in 96 well plates. Single clones were confirmed for integration of MYC tag at C-terminus (Fig. S2).

### Western blotting

Parasites growing in confluent HFF cells were mechanically lysed and filtered to remove host cell debris followed by centrifugation. Parasite pellets were lysed in NuPAGE buffer, sonicated, and boiled for 5 minutes. Parasite lysates were centrifuged briefly to remove debris and separated on 4-12% NuPAGE gels (Invitrogen). Separated proteins were transferred to PVDF membrane and blocked in 5% Nonfat dry milk-TBST solution for 1 hour at room temperature. Blocked PVDF membranes were incubated with anti-HA (1:2000, Roche), anti-MYC (1:3000, Cell Signaling) or anti-SAG1 (1:5000, ThermoFisher) antibodies in blocking buffer at 4°C overnight with gentle shaking. Membranes were washed with TBST and incubated with HRP-conjugated anti-rat or anti-mouse (1:2000; GE healthcare) for 1 hour at room temperature. After washing with: TBST, membranes were developed with SuperSignal West Femto substrate (ThermoFisher).

### Immunofluorescence assay

Parasites growing in HFF monolayers were fixed in 4% paraformaldehyde solution for 15 minutes at room temperature followed by washing with phosphate buffered saline (PBS). Fixed cells were permeabilized and blocked with blocking buffer (3% BSA + 0.2% Triton X-100 in PBS) for 1 h. Cells were incubated with primary antibodies in blocking buffer at 4°C temperature overnight. Cells were washed with PBS and finally incubated with DAPI (1:1000) and Alexa fluor secondary antibodies coupled with desired fluorophore for 1 hour at room temperature. Cells were washed and mounted with Prolong Gold anti-fade reagent (Invitrogen) before analysis with a Leica DMI6000 B inverted microscope. Dilutions for antibodies used for IFA are as follows: Rabbit α-HA (1:2000, Sigma, H6908), Mouse mAb α-MYC (1:1000, Cell signaling, 9B11), Rabbit α-MYC (1:2000, ThermoFisher, PA1-981) and Mouse mAb 5F4 (1:5000, F1B ATPase, a gift from Peter Bradley), Alexa Fluor 488/594 secondary antibodies (1:2000, ThermoFisher) were used as appropriate.

### RNA sequencing

Parasites were allowed to invade confluent HFF monolayers for 24 h. Infected HFF monolayers were scraped, and syringe lysed with 23-gauge needle to release intracellular parasites and passed through a 3 μM filter. Filtered parasites were centrifuged at 300 x *g* for 15 minutes and DMEM media was removed to isolate the parasite pellet. The parasite pellet was resuspended in 1 x PBS and centrifuged at 300 x g for 15 minutes, twice. Total RNA was extracted with TRIzol reagent (Invitrogen) as per manufacturer’s protocol and measured by Nanodrop. RNAseq libraries were prepared from polyA-purified mRNA by Azenta using standard Illumina protocols. Libraries were sequenced PE x 150bp.

### RNAseq data analysis

RNAseq read data generated from either the RH or the ME49 genetic background were aligned to the ME49 reference sequence (v52) with hisat2 using default settings (38). Differential gene expression was conducted with DESeq2 using default settings (39). Raw sequencing data and processed files generated in this study are available at GEO GSE217226 and data pertaining to HDAC3 and MORC impairment (6) were obtained from GEO GSE136123.

### CUT&Tag assay

Chromatin occupancy for ME49^TIR1^ parasites expressing AP2XII-2^AID-HA^ was performed by CUT&Tag profiling (14). Briefly, 2×10^7^ intracellular tachyzoites were syringe lysed 24 hours post invasion to release fresh parasites. Parasites were centrifuged at 600 x *g* for 10 minutes at room temperature. Parasite pellets were then processed as per the manufacturer’s protocol (CUT&Tag-IT™ Assay Kit, Anti-Rabbit, Active motif, Cat no. 53160). Libraries were sequenced by Azenta at PE 2×150 bp.

### Chromatin occupancy data analysis

Sequencing data was aligned to the ME49 genome (v52) with bowtie2 (40) allowing for mate dovetailing. Peaks were called with MACS2 using default settings (41). Peak annotation was performed with Homer using a 2kb cutoff distance (42). Peak overlap was conducted with bedtools overlap function (43). AP2XII-2 data was deposited at GEO GSE217220 and raw data pertaining to ChIPseq of MORC and HDAC3 (6) were obtained from GEO GSE136060.

## ACKNOWLEDGEMENTS

The authors would like to thank Dr. Peter Bradley for the F_1_B ATPase antibody, Dr. Vicki Jeffers for technical advice related to CUT&Tag, and members of the Sullivan lab and Biology of Intracellular Pathogens group at the IU School of Medicine for helpful discussions. This research was supported by National Institutes of Health grants AI124682 (M.W.W.) and AI152583 (W.J.S.). The funders had no role in study design, data collection and interpretation, or the decision to submit the work for publication.

## SUPPLEMENTAL FIGURE LEGENDS

**Figure S1. Re-analysis of transcriptomic changes that occur upon impairment of HDAC3/MORC complex.** A-B) Differential gene expression analysis for HDAC3 inhibition (A) and IAA-induced MORC depletion (B), each conducted as it was for newly generated AP2XII-2 depletion data.

**Figure S2. Validation for endogenous tagging of AAH1 and AAH2.** A) AAH1 was endogenously tagged with a MYC epitope tag using a CAS9-assisted approach to facility double homologous recombination as indicated. B) AAH2 was endogenously tagged with a MYC epitope tag using a single crossover approach as indicated. C-D) Diagnostic PCRs checking for correct integration (I) or the unmodified endogenous locus (E) were performed on parental and tagged lines. Specific regions selected for PCR analysis are highlighted in (A) and (B).

## SUPPLEMENTAL TABLE LEGENDS

**Table S1. Chromatin-associated regions for AP2XII-2, HDAC3, and MORC as revealed by CUT&Tag or ChIPseq.**

**Table S2. Differential gene expression analysis resulting from AP2XII-2 depletion in RH and ME49 backgrounds.**

**Table S3. Differential gene expression analysis resulting from HDAC3 inhibition or MORC depletion.**

**Table S4. List of primers used in this study.**

## Notes

### Competing Interest Statement

The authors have declared no competing interest.

### Summary of Updates

Supplemental files were uploaded.

