## Supplementary figures and images for "AP2XII-2 coordinates transcriptional repression for *Toxoplasma gondii* sexual commitment"

### Fig. S1

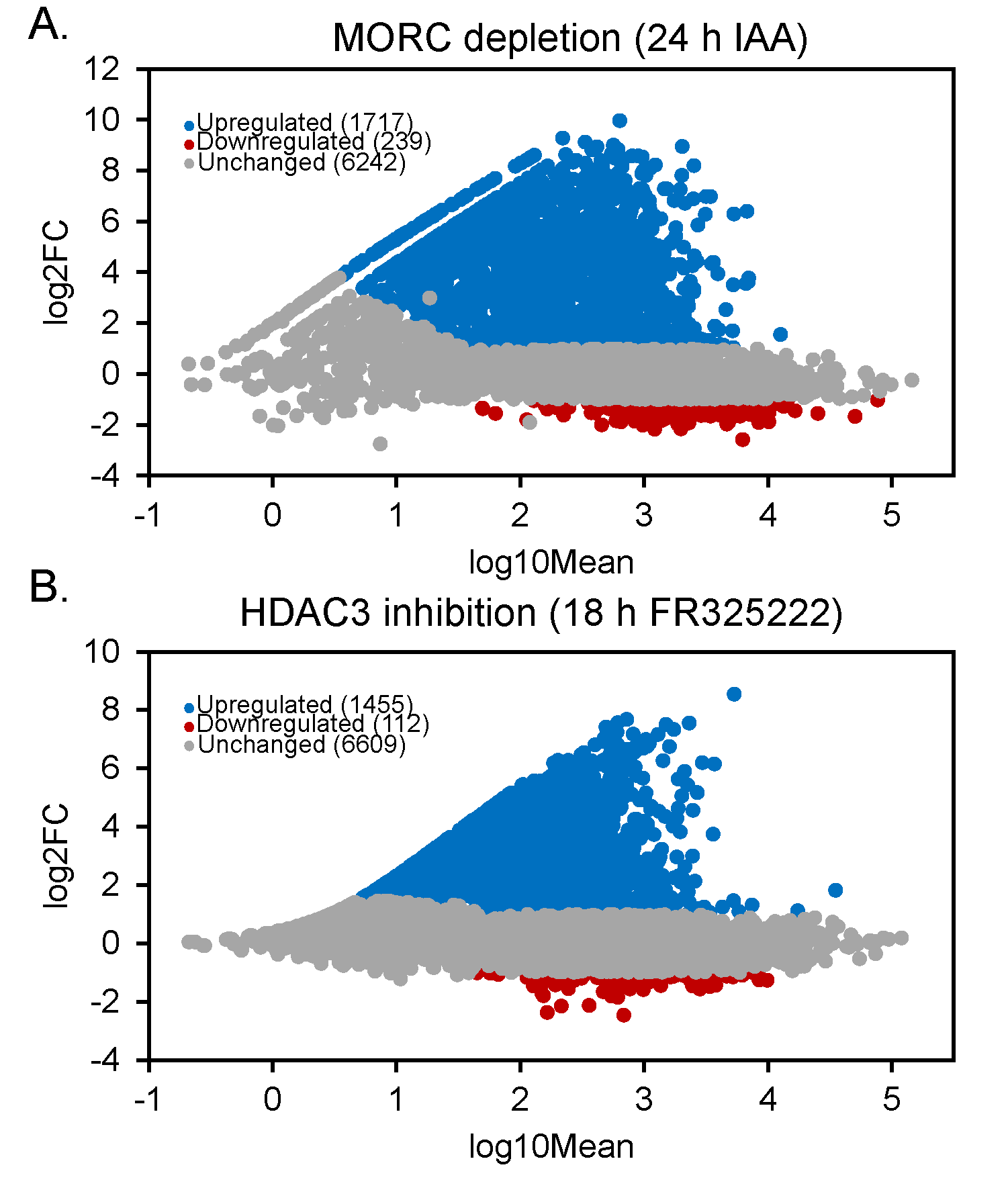

### Fig. S2

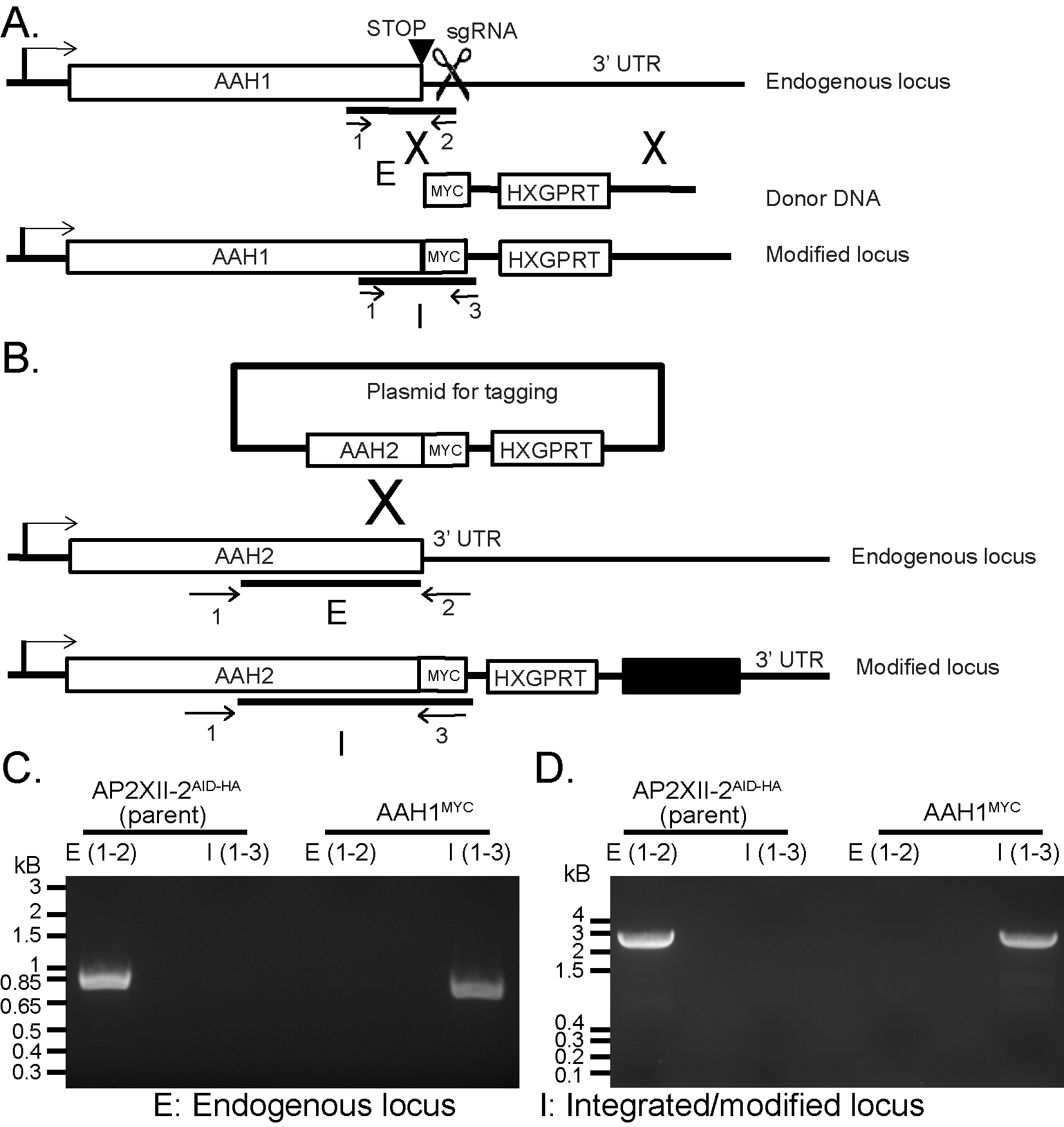
